# Optical topometry and machine learning to rapidly phenotype stomatal patterning traits for QTL mapping in maize

**DOI:** 10.1101/2020.10.09.333880

**Authors:** Jiayang Xie, Dustin Mayfield-Jones, Gorka Erice, Min Choi, Andrew D.B. Leakey

## Abstract

Stomata are adjustable pores on leaf surfaces that regulate the trade-off of CO_2_ uptake with water vapor loss, thus having critical roles in controlling photosynthetic carbon gain and plant water use. The lack of easy, rapid methods for phenotyping epidermal cell traits have limited the use of quantitative, forward and reverse genetics to discover the genetic basis of stomatal patterning. A new high-throughput epidermal cell phenotyping pipeline is presented here and used for quantitative trait loci (QTL) mapping in field-grown maize. The locations and sizes of stomatal complexes and pavement cells on images acquired by an optical topometer from mature leaves were automatically determined. Computer estimated stomatal complex density (SCD; R^2^ = 0.97) and stomatal complex area (SCA; R^2^ = 0.71) were strongly correlated with human measurements. Leaf gas exchange traits correlated with the dimensions and proportion of stomatal complexes but, unexpectedly, did not correlate with SCD. Genetic variation in epidermal traits were consistent across two field seasons. Out of 143 QTLs in total, 36 QTLs were consistently identified for a given trait in both years. 24 hotspots of overlapping QTLs for multiple traits were identified. Orthologs of genes known to regulate stomatal patterning in *Arabidopsis* were located within some, but not all, of these regions. This study demonstrates how discovery of the genetic basis for stomatal patterning can be accelerated in maize, a model for C_4_ species where these processes are poorly understood.

**One sentence summary:** Optical topometry and machine learning tools were developed to assess epidermal cell patterning, and applied to analyze its genetic architecture alongside leaf photosynthetic gas exchange in maize.

## INTRODUCTION

Stomata are the adjustable pores on leaf surfaces that regulate gas exchange, most notably CO_2_ uptake and water vapor loss. The ratio of carbon gained to water lost is defined as water use efficiency (WUE), and represents arguably the most fundamental trade-off faced by land plants (Leakey et al., 2019). The pattern of stomata on the epidermis, and the dynamics of stomatal opening and closing, influence many important processes from food and energy production to global carbon and water cycling (Hetherington and Woodward, 2003). The accessibility of stomata on the plant exterior surface has also made them a model system for studying developmental and signaling processes (Blatt, 2000; Schroeder et al., 2001; Bergmann, 2004; Lawson et al., 2014; Torii, 2015). Consequently, there is significant potential for fundamental scientific discoveries about stomata to be leveraged for improvement of crop performance and sustainability through breeding or biotechnology (Yoo et al., 2010; Franks et al., 2015; Hughes et al., 2017; Caine et al., 2019; Lawson and Vialet-Chabrand, 2019; Harrison et al., 2020; McKown and Bergmann, 2020).

Despite the accessibility and importance of stomata, assessing the patterning of epidermal cells has remained a laborious and time-consuming task for many decades. Most studies of stomatal patterning still rely on methods of imprinting or peeling the epidermis from live tissue, followed by light microscopy, and manual identification and measurement of cells in images (e.g. Biscoe, 1872; Caine et al., 2019; Vőfély et al., 2019). This limits the application of quantitative, forward and reverse genetics to understand the genes and processes that regulate stomatal patterning. And, it means samples cannot be analyzed with sufficient throughput for stomatal patterning to be a target trait in modern crop breeding programs.

Optical topometry (OT) is a rare example of a new methodology proposed to accelerate the acquisition of epidermal patterning data through rapid image acquisition. OT is a non-destructive method for use on fresh or frozen leaf samples, which requires no sample preparation beyond sticking a piece of leaf to a microscope slide with double-sided sticky tape (Haus et al., 2015). It gathers focused pixels across plains of the leaf surface in less than one minute per field of view. OT images have been manually counted to assess stomatal density responses to elevated [CO_2_] in *Arabidopsis* (Haus et al., 2018). But, an automated analysis pipeline is still needed to robustly capture within-species genetic variation in epidermal patterning from OT images with the fidelity required for genetic analysis.

There have been many attempts to address the phenotyping bottleneck for stomatal patterning through computer-aided image analysis. Classical image processing methods (Omasa and Onoe, 1984; Liu et al., 2016; Duarte et al., 2017) and machine learning models have been applied (Vialet-Chabrand and Brendel, 2014; Higaki et al., 2015; Jayakody et al., 2017; Saponaro et al., 2017; Dittberner et al., 2018; Toda et al., 2018; Bhugra et al., 2019; Sakoda et al., 2019; Aono et al., 2019; Fetter et al., 2019; Li et al., 2019). While a number of these methods have been demonstrated to work within constrained image sets, none of them have been widely adopted, even within a single species. Some of these tools require scanning electron microscopy (SEM), adding to the sample preparation and image acquisition burden (Aono et al., 2019; Bhugra et al., 2019; Fetter et al., 2019). Most existing tools are limited to identifying and phenotyping stomatal complexes. Adding the ability to measure pavement cells is valuable in its own right and also allows calculation of stomatal index (SI; i.e. the ratio of stomata number to all epidermal cell number given in unit leaf area). SI is a key trait because it is directly influenced by mechanisms that regulate epidermal cell fate and it is less sensitive to environmental influences than stomatal density (Royer, 2001). Therefore, developing an end-to-end pipeline for rapid acquisition and comprehensive analysis of epidermal cell patterning, and demonstrating its application in investigation of genetic variation in stomatal patterning, remains an important but elusive goal.

In recent years, important progress has been made in studying the degree to which orthologs of stomatal patterning genes in *Arabidopsis* (Pillitteri and Torii, 2012) have conserved or novel functions in C_3_ grass species (Raissig et al., 2016; Hughes et al., 2017; Raissig et al., 2017; Yin et al., 2017; Hepworth et al., 2018; McKown and Bergmann, 2020). But, very little is known about the trait relationships and genetic control of stomatal patterning and iWUE in C_4_ species (Leakey et al., 2019). And, apart from a few notable examples (Cartwright et al., 2009; Campitelli et al., 2016; Raissig et al., 2017) quantitative genetics and forward genetic screens to identify putative regulators of stomatal patterning still have generally not met their potential to drive discovery of genotype-to-phenotype relationships.

Linkage mapping in barley, wheat, and rice has discovered QTLs that are associated with stomatal patterning traits (Patto et al., 2003; Laza et al., 2010; Liu et al., 2014; Liu et al., 2017; Sumathi et al., 2018), including some that co-localize with yield QTL (Shahinnia et al., 2016). But, the only reports of similar experiments in maize predate statistical techniques such as QTL mapping (Heichel, 1971). Maize is the most important crop in the world in terms of total production (USDA, 2019), with the Midwest U.S. producing approximately 27% of the global harvest (USDA-FAS, 2020). Maize yield is limited by water availability, and increasingly sensitive to drought as a side effect of productivity increases resulting from improved breeding and management (Lobell et al., 2014). Conversely, increased maize production over recent decades has led to faster water cycling and regional cooling in Midwest U.S. (Alter et al., 2018). Therefore, improved understanding of the genetic basis for variation in stomatal traits in maize has implications for agricultural productivity, resilience and sustainability. And, maize is a highly tractable, model experimental system for crop genetics (Buckler et al., 2009).

In summary, the current study was motivated by the need for a tool to accelerate phenotyping of epidermal cell patterning, which could then be demonstrated by applying it to investigate the genetic architecture of stomatal patterning traits in maize. The desired characteristics of an end-to-end phenotyping pipeline are: (1) little to no sample preparation and quick image acquisition; (2) fast, accurate and robust detection of epidermal cells; and (3) the ability to extract the number, size and position of pavement cells as well as stomatal complexes. OT was tested as a data acquisition method from leaves that were stored frozen after being grown in the field. For epidermal cell detection, the recently developed Mask R-CNN model for object instance detection (He et al., 2017) was tested to treat stomata and pavement cells as two object classes, so that their position and size could be extracted simultaneously. A RIL population resulting from a B73 x MS71 cross was grown in two years in the field. Stomatal patterning was phenotyped along with leaf photosynthetic gas exchange and specific leaf area to investigate the genetic architecture of these important traits in a major crop and model C_4_ species.

## RESULTS

### High throughput phenotyping pipeline for epidermal cells of maize

A high throughput epidermal cell detection pipeline requires both efficient image acquisition and automatic cell detection (Fig. 1). Optical topometry (OT) allowed rapid, nondestructive imaging of leaf samples. Less than 1 minute was required from locating the portion of epidermis to be scanned to outputting a 3D topography surface layer with dimensions of 0.8mm x 0.8mm (e.g. Fig. 2A). Overall, 7033 fields of view were sampled from 1569 leaf samples collected over two field seasons, with scanning being completed in approximately 24 person days. The Mask-RCNN model automatically detected stomatal complexes as well as pavement cells, even though the latter varied greatly in their physical shape and size (Fig. 2). Analysis of a full image set for QTL mapping (~4000 images) was completed in approximately 120 h (Table 1).

**Table 1.**
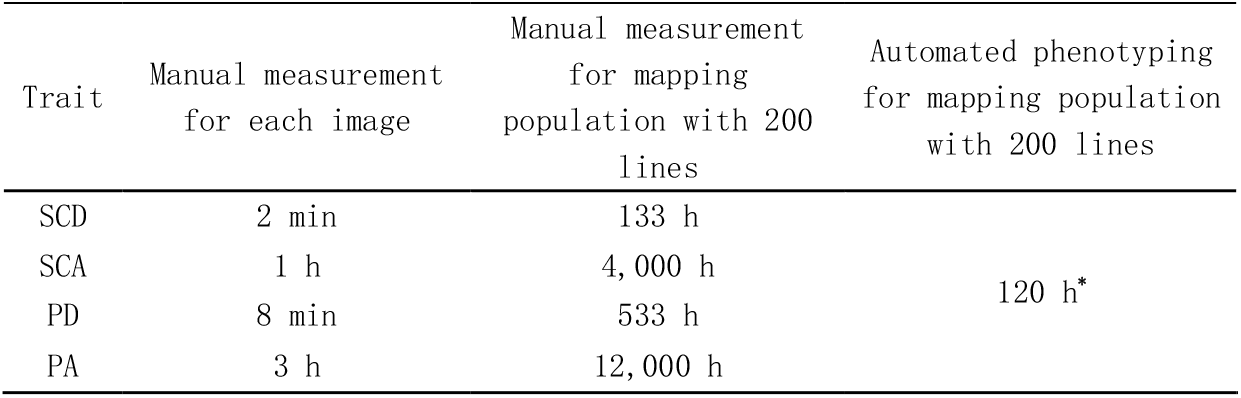
Time investment approximations for epidermal cell detection and trait extractions comparing manual measurements versus automated detections. SCD, Stomatal complex density; SCA, stomatal complex area; PD, pavement cell density; PCA, pavement cell area; h, hours. Estimations were done on 20X magnification maize abaxial images (0.8mm x 0.8mm) for a mapping population with 200 lines, 4 replications and 5 leaf level sub-samples (4000 images). Asterisk designates time estimation for all traits combined.

**FIGURE 1.**
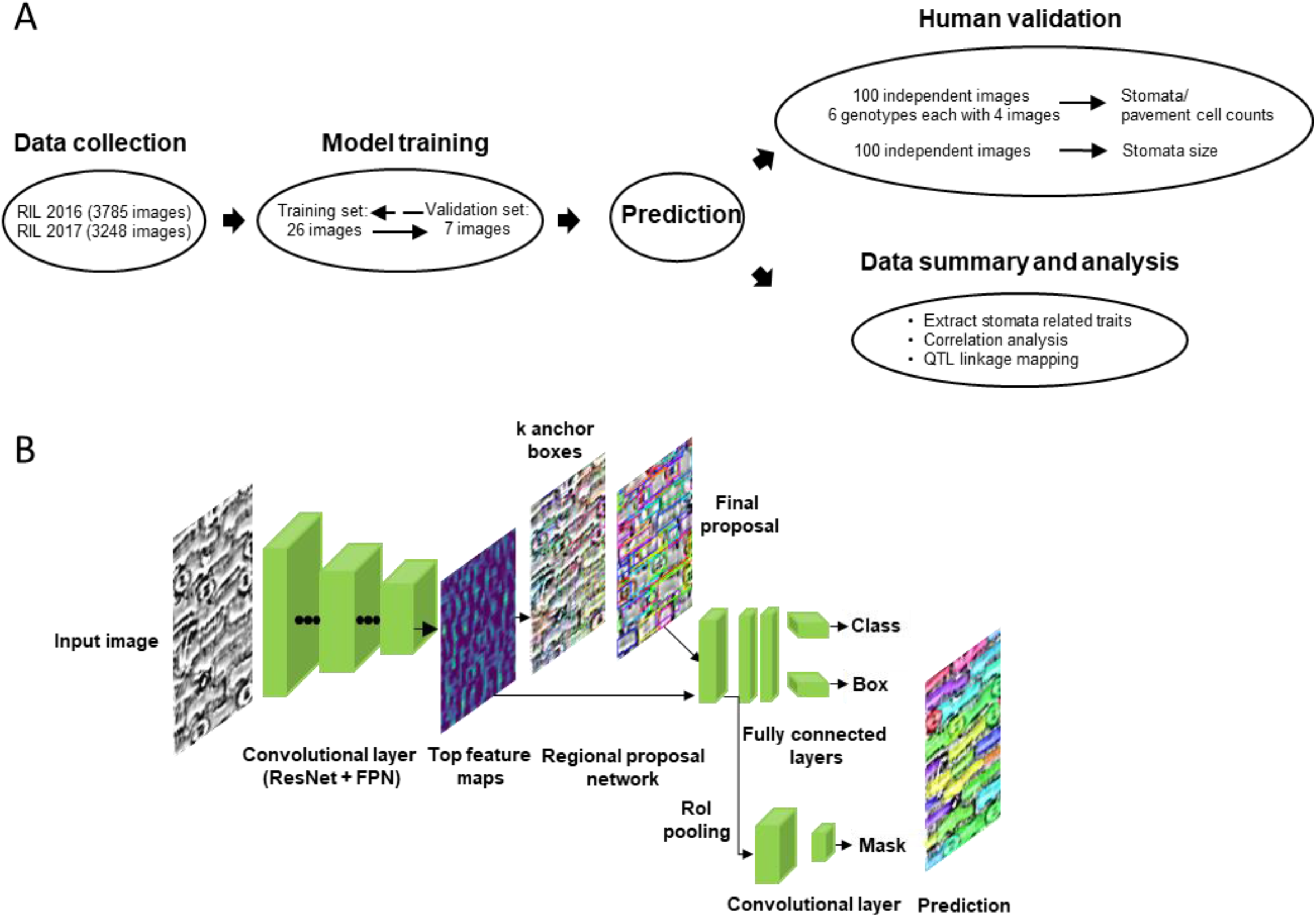
Workflow of data collection, model training, model prediction, human validation and experimental data analysis used to phenotype epidermal cell patterning traits (A). Summary of pipeline used by Mask R-CNN to analyze images captured by optical tomography for stomata and pavement cell detection. Image example was truncated from standard image.

**FIGURE 2.**
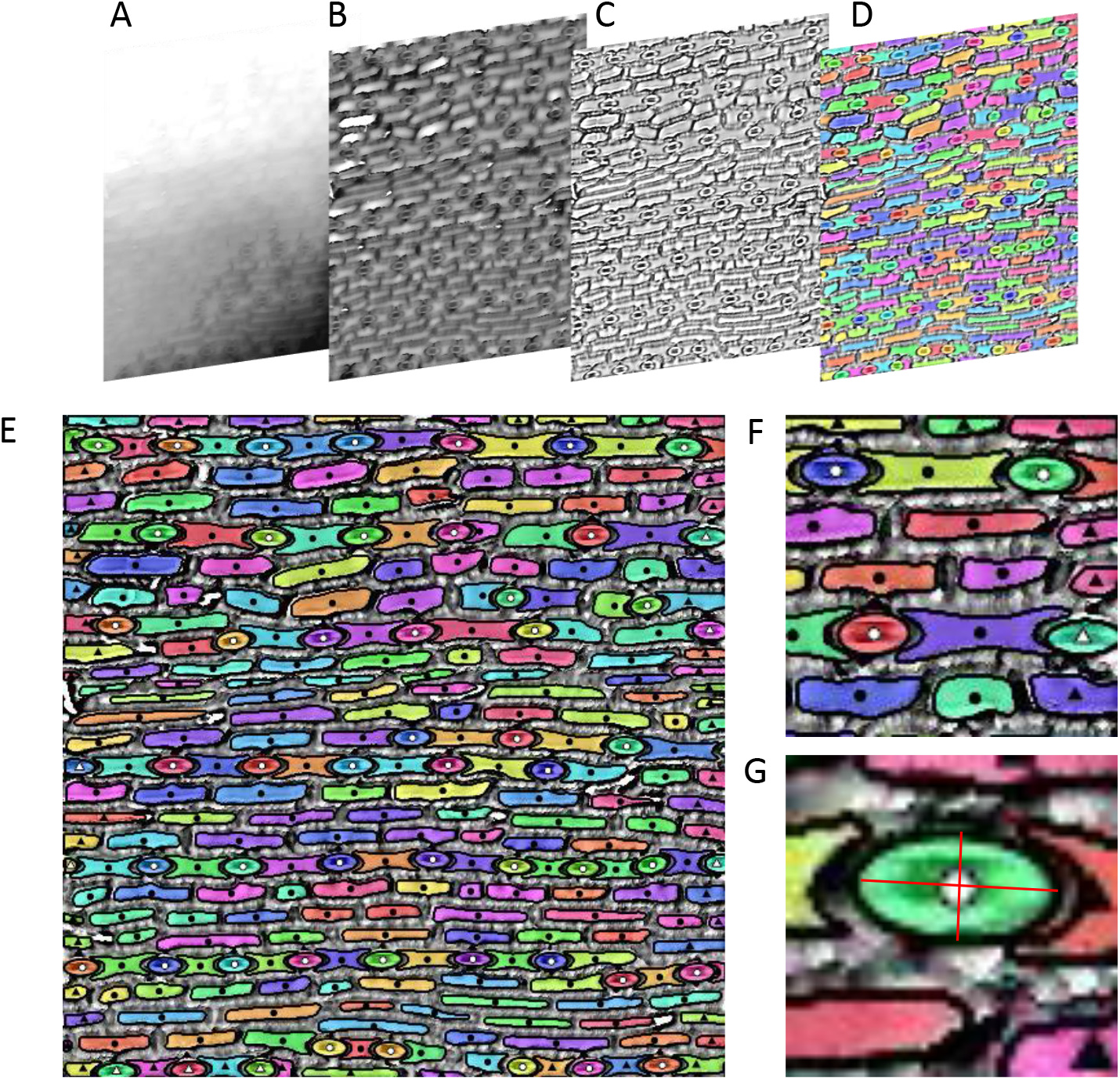
Example steps in the process of analyzing an optical tomography image for epidermal cell patterning, including: the 3D topography image layer extracted from raw filers output by the optical topometer (A); flattening by use of Robust Gaussian filters (B); contrast enhancement by use of a Laplacian filter (C); prediction of cell instances by Mask R-CNN (D, E, F, G). Cell related traits were calculated and extracted based on cell boundary coordinates, with boundary and centroid labeled for better visualization (E). Zooming in shows stomata were labeled with white centroids while pavement cells were labeled with black centroids (F). Cells that were cut off on image edges were tagged with triangles and were excluded in estimation of average cell size. Ellipses were fit to stomatal complexes, with width and length calculated as the lengths of minor and major axis of the ellipse (red lines; G).

### Human validation of MASK R-CNN cell counts and stomatal complex size

Variation among 6 trained human evaluators contributed a small portion of the variance within the dataset for both SCD (2%) and pavement cell density (PD; 6%) (Fig. S3). Variation among evaluators contributed a greater proportion of variance for stomatal complex width (56 %), stomatal complex length (23 %) and stomatal complex area (15 %). Nonetheless, uncertainty around the mean value of human measurements was low (expressed as standard error around plotted data in Fig. 3, A and B). There was no variance in estimates of cell density from Mask R-CNN when the same image was repeatedly submitted to the analysis pipeline, so no measure of technical variation could be expressed.

**FIGURE 3.**
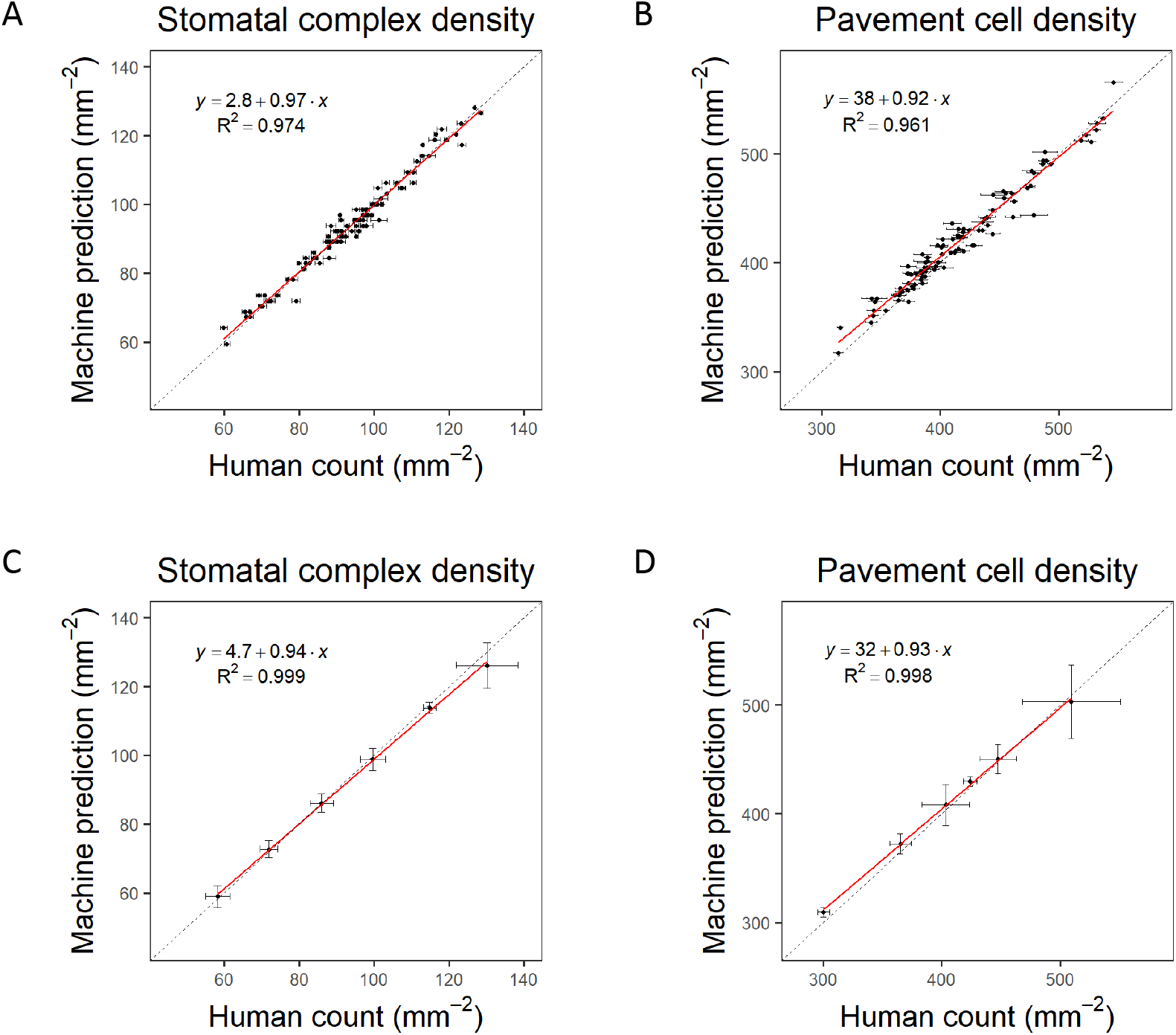
Scatterplots of stomatal patterning traits comparing data measured by humans versus data measured by the computer using MASK R-CNN: stomatal complex density (A,C); and pavement cell density (B,D). Plotted data describe 100 randomly selected optical tomography images from the B73 x MS71 maize RIL population with error bars showing the standard error of technical variation among six expert human evaluators on each individual image (A,B) or genotype means for 6 RILs selected to represent the range of observed trait values in the population with error bars showing the standard error of biological variation among replicates based on the mean of predictions from six expert human evaluators or computer predictions using MASK R-CNN (C,D). There is no variance among predictions by MASK R-CNN when it is presented with a given image multiple times. The line of best fit (red line) and 1:1 line (black dashed line) are shown along with the correlation coefficient (r^2^).

The mean density of cells estimated by the group of human evaluators was very strongly correlated with computer estimation of both SCD (R^2^ = 0.97, p<0.0001; Fig. 3A) and PD (R^2^ = 0.96, p<0.0001; Fig. 3B) and displayed very low bias from the 1:1 line. The mean data from human evaluators were also highly significantly correlated with computer measurements for stomatal complex length (SCL; R^2^ = 0.81, p<0.0001; Fig. 4A), stomatal complex width (SCW; R^2^ = 0.54, p<0.0001; Fig. 4B) and stomatal complex area (SCA; R^2^ = 0.71, p<0.0001; Fig. 4C). All three traits were slightly underestimated by machine measurements relative to human measurements, with the absolute bias being greater for larger cells than small cells.

**FIGURE 4.**
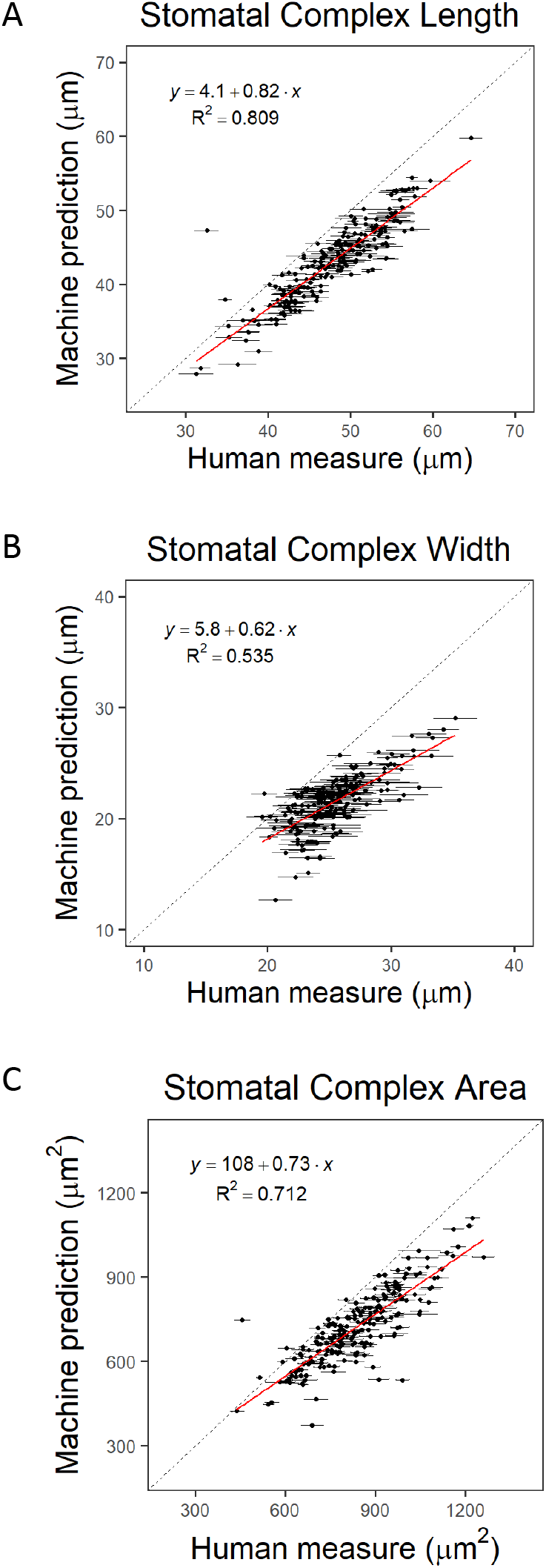
Scatterplots of stomatal complex length (A), stomatal complex width (B) and stomatal complex area (C) comparing data measured by humans versus data measured by the computer using MASK R-CNN: Plotted data describe 210 stomatal complexes (5 each from 42 images) randomly selected from the B73 x MS71 maize RIL population with error bars showing the standard error of technical variation among six expert human evaluators on each individual image. There is no variance among predictions by MASK R-CNN when it is presented with a given image multiple times. The line of best fit (red line) and 1:1 line (black dashed line) are shown along with the correlation coefficient (r^2^).

To further evaluate sources of variation in stomatal patterning traits, six RILs were chosen that represented the range of SCD observed across the full mapping population in the 2016 growing season. All the images for those six RILs were then manually counted by five human beings as well as by machine. Variation around the genotype means derived from machine counts was similar or smaller than the variation resulting from using the mean of five expert evaluators as the input (expressed as standard error around plotted data in Fig. 3, C and D). Genotype mean values based on machine counts were very strongly correlated with best-estimates from human evaluators for both stomatal complex density (R^2^ = 0.999, p<0.0001; Fig. 3C) and pavement cell density (R^2^ = 0.998, p<0.0001; Fig. 3D), and had very little bias from the 1:1 line.

### Genotypic variation in traits within and across years

Genotypic variation in stomatal patterning traits displayed good repeatability across growing seasons (Fig. 5). Genotype means were significantly correlated across the two years for all traits assessed with goodness-of-fit (R^2^) ranked from highest to lowest of: 0.70 for SCTA; 0.69 for SPI, 0.68 for SI, 0.64 for SCD; 0.64 PA; 0.60 for PD; 0.56 for SCL; 0.52 for SCLWR; 0.50 for SCA; 0.46 for SCW; 0.43 for PTA; and 0.13 for SLA.

**FIGURE 5.**
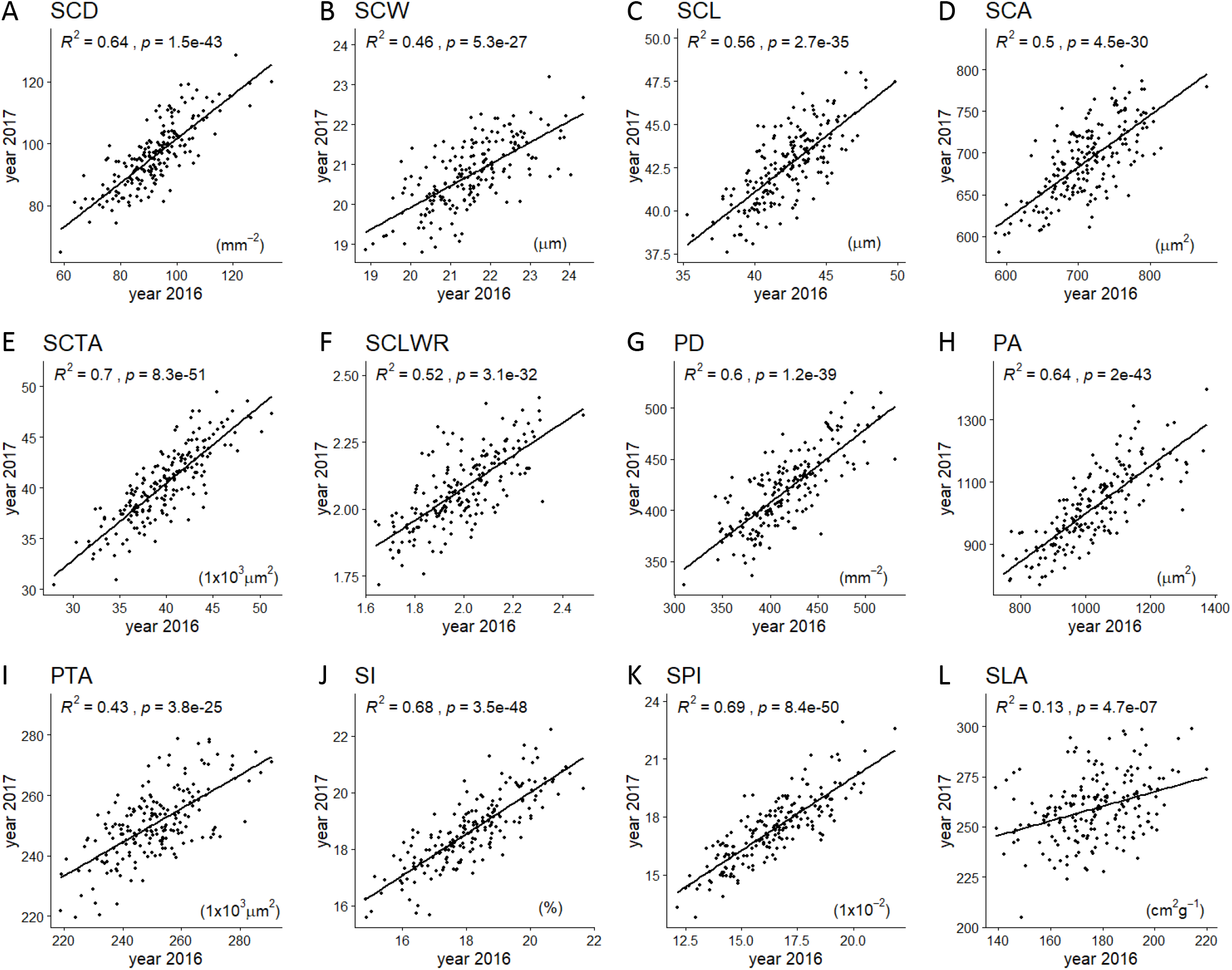
Scatterplots of stomatal complex density (SCD, A), stomatal complex width (SCW, B), stomatal complex length (SCL, C), stomatal complex area (SCA, D), stomatal complex total area (SCTA, E), stomatal complex length to width ratio (SCLWR, F), pavement cell density (PD, G), pavement cell area (PA, H), pavement cell total area (PTA, I), stomatal index (SI, J), stomatal pore area index (SPI, K), specific leaf area (SLA, L) comparing genotype means for 191 maize B73 x MS71 RILs grown during the 2016 versus 2017 field seasons. The line of best fit (black line), correlation coefficient (r^2^) and associated p-value are shown.

Among the 198 RILs assessed over the two years, the relative range of stomatal patterning traits varied from more than 2-fold, i.e., 127% for SCD (59 to 134 mm^−2^) down to 29% for SCW (18.8 to 24.3 μm; Fig. S4). Specific leaf area (SLA) was significantly greater in 2017 (205 to 299 cm^2^g^−1^) compared to 2016 (139 to 220 cm^2^g^−1^). In 2017, leaf photosynthetic gas exchange traits varied 2-4 fold among the 192 RILs for the rate of CO_2_ assimilation (*A*), stomatal conductance (*g_s_*); the ratio of intercellular [CO_2_] to atmospheric [CO_2_] (*c_i_/c_a_*); and intrinsic water use efficiency (*iWUE*). The ranges of all trait values significantly exceeded the trait variation between the parent lines B73 and MS71 (Fig. S4). As expected, SCD and SI were significant lower in MS71 than B73. This corresponded with greater stomatal complex size in MS71 compared to B73 in terms of SCW, SCL and SCA. SCLWR was greater in MS71 than B73. In terms of leaf gas exchange, MS71 had lower *g_s_,* lower *A*, lower *C_i_/c_a_* and greater *iWUE* than B73 (Fig. S4).

### Trait relationships

Correlation matrices for genotype means of stomatal patterning traits were very similar for data collected in 2016 (Fig. S5) and 2017 (Fig. 6). Therefore, the presentation of results here will focus on data from 2017, when anatomical traits were assessed alongside leaf photosynthetic gas exchange.

**FIGURE 6.**
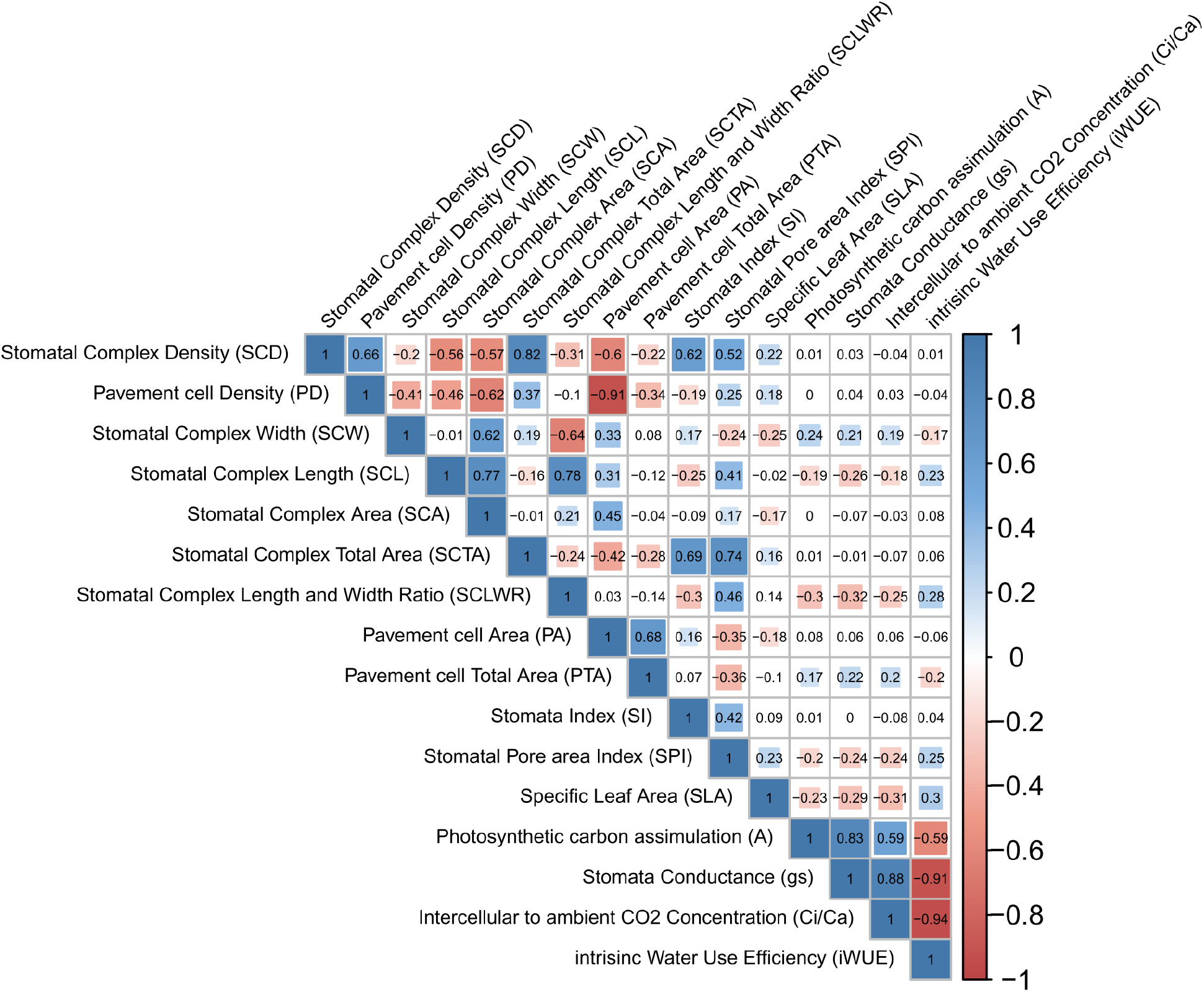
Correlation matrix for stomatal complex density (SCD), stomatal complex width (SCW), stomatal complex length (SCL), stomatal complex area (SCA), stomatal complex total area (SCTA), stomatal complex length to width ratio (SCLWR), pavement cell density (PD), pavement cell area (PA), pavement cell total area (PTA), stomatal index (SI), stomatal pore area index (SPI), specific leaf area (SLA), rate of photosynthetic CO_2_ assimilation (*A*), stomatal conductance (*g_s_*), ratio of leaf intercellular to atmospheric CO_2_ concentration (*c_i_/c_a_*) and intrinsic water use efficiency (*iWUE*), based on genotype means of the maize B73 x MS71 RIL population grown in 2017 (n = 194). Statistically significant correlations (p<0.05) are highlighted with colored cells that reflect the strength of the correlation by the size of the shaded area and are colored from red (positive correlation, coefficient = 1) to blue (negative correlation, coefficient = −1).

There were numerous significant trait associations among anatomical stomatal patterning traits and also among leaf photosynthetic gas exchange traits. Genotypes with larger stomatal complexes tended to have larger pavement cells (SCA vs PA, r = 0.45), which resulted in a positive correlation between SCD and PD as well (r = 0.66). SCD was negatively correlated with measures of stomatal complex size, including SCW (r = −0.2), SCL (r = −0.56) and SCA (r = −0.57). As SCD increased it was associated with a significant decrease in SCLWR (i.e., rounder or less elongated stomatal complexes, r = −0.31). But, PD was not significantly correlated with the shape of stomatal complexes, SCLWR (p = 0.16). With the majority of the epidermis occupied by pavement cells, the trade-off between density (PD) and size (PA) was even stronger than for stomatal complexes (r = −0.91). After aggregating across the epidermis, SCTA was positively correlated with SCD (r = 0.82) and SI (r = 0.69) but was influenced in a mixed and weaker manner by stomatal complex size or proportions in terms of SCW (r = 0.19), SCL (r = −0.16), SCA (p = 0.88) or SCLWR (r = −0.24). Considering just cell identity, SI was more strongly correlated with variation in SCD (r = 0.62) than PD (r = −0.19). Meanwhile, there were strong positive correlations of *g_s_* with *A* (r = 0.83) and *gs* with *c_i_/c_a_* (r = 0.88). And a correspondingly strong negative correlation of *g_s_* with *iWUE* and (r = −0.91). There were weaker, but significant correlations between *A* and *c_i_/c_a_* (r = 0.59) and *A* and *iWUE* (r = −0.59). SLA was positively correlated with *iWUE* (r = 0.30) while being negatively correlated with *A* (r = −0.23), *g_s_* (r = −0.29) and *c_i_/c_a_* (r = −0.31).

Examining structure-function relationships across trait categories, *A*, *g_s_*, *c_i_/c_a_* and *iWUE* were not significantly correlated with measures linked to the number or overall size of stomatal complexes (i.e. SCD, SCA or SCTA). However, traits including the component dimensions of stomatal complexes (i.e. SCL, SCLWR, and SPI) were negatively correlated with *A*, *g_s_*, and *c_i_/c_a_* and positively correlated with *iWUE*. And, SCW was positively correlated with *A*, *g_s_*, and *c_i_/c_a_* and negatively correlated with *iWUE*.

### Linkage mapping

143 individual QTL were identified (Fig. 7, Table S1) in total for the 16 traits tested in 2016 (60 QTL) and 2017 (83 QTL). Almost half of these QTL were independently identified for the same trait in both years, providing greater confidence for significant genotype to phenotyping associations at 36 loci spread across every chromosome except chromosome 4. The percentage of phenotypic variance explained (PVE) by individual QTL was 8.2 % on average, with a maximum of 18.3 % for PA at Chr9A (Fig. 7, Table S1). For the anatomical stomatal patterning traits tested in both years, the number of QTL identified varied from five QTL for SCL and six QTL for SPI to 18 QTL for SI and 20 QTL for SCD (Fig. 7, Table S1). In comparison, one to three QTL were identified for each of the functional leaf photosynthetic gas exchange traits, which were only tested in 2017. Correspondingly, the total PVE by all the QTL for a given trait was greater for the anatomical stomatal patterning traits (51 % on average in 2017) than for the photosynthetic gas exchange traits (17 % on average in 2017; Fig. S6). In addition, for the anatomical stomatal patterning traits, the total PVE was generally equivalent or greater in 2017 (51 % on average) than in 2016 (45 % on average, Fig. S6). The traits with the greatest total PVE (i.e. > 50%) were SI, SCA, SCD, SCTA and PA, although total PVE was >35 % for all anatomical traits.

**FIGURE 7.**
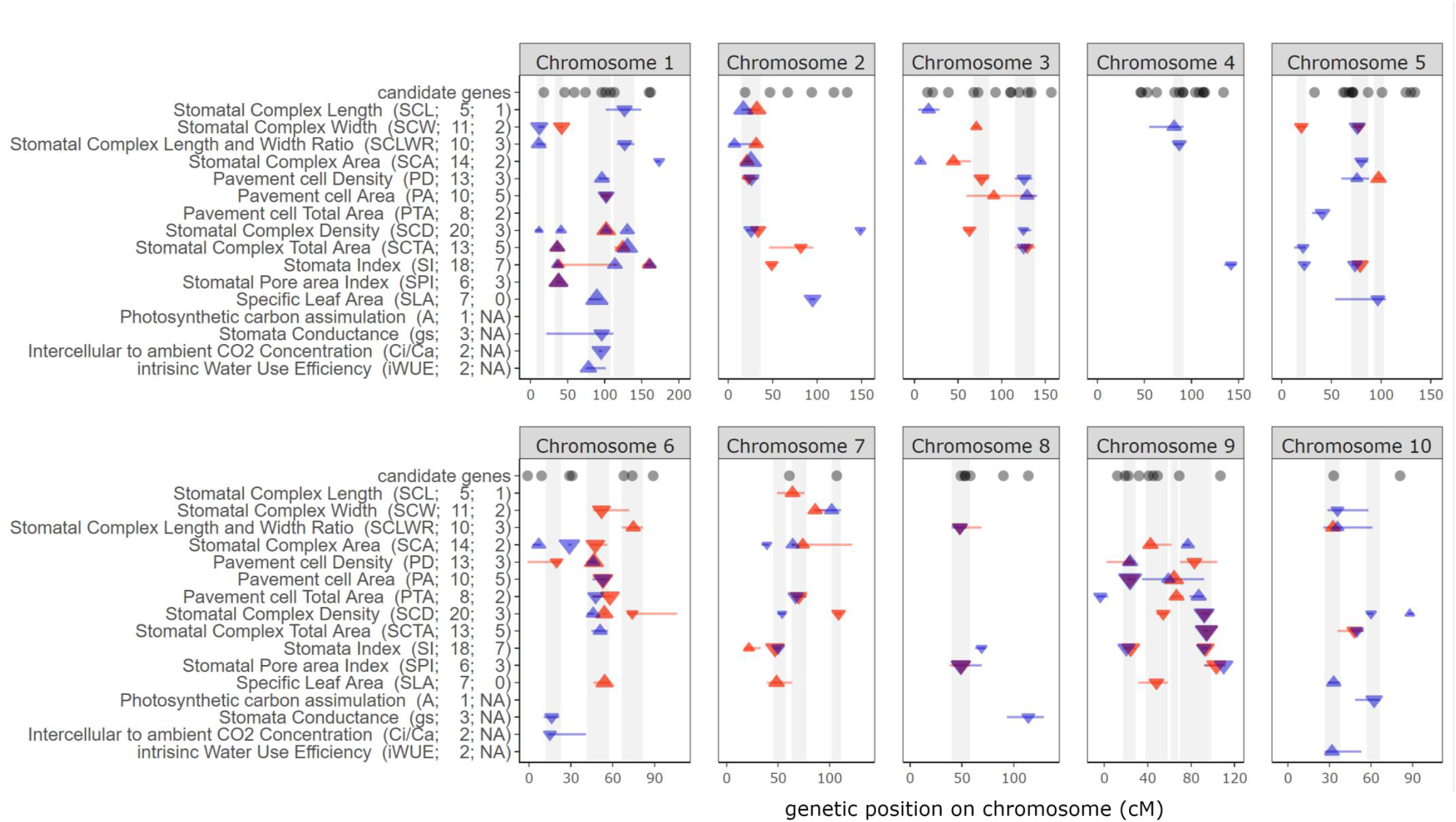
QTL mapping for stomatal complex density (SCD), stomatal complex width (SCW), stomatal complex length (SCL), stomatal complex area (SCA), stomatal complex total area (SCTA), stomatal complex length to width ratio (SCLWR), pavement cell density (PD), pavement cell area (PA), pavement cell total area (PTA), stomatal index (SI), stomatal pore area index (SPI), specific leaf area (SLA), rate of photosynthetic CO_2_ assimilation (*A*), stomatal conductance (*g_s_*), ratio of leaf intercellular to atmospheric CO_2_ concentration (*c_i_/c_a_*) and intrinsic water use efficiency (*iWUE*) from the B73 x MS71 RIL population. Each panel corresponds to an individual chromosome, where the values on the x-axis are chromosome position (cM). Numbers in parentheses following abbreviated trait names on the y-axis indicate the total number of QTL for that trait detected across the two growing seasons and the number of QTL for that trait that were detected consistently across both growing seasons. Each triangle represents a single QTL detected, with the direction of the arrow corresponding to the directional effect of the MS71 allele. Triangles are colored to indicate QTLs that were significant in 2016 (red), 2017 (blue), or overlapping across both years (purple). Error bars indicate the 1.5 LOD support intervals. Grey shaded areas indicate clusters of co-located QTL. The location of orthologs of known stomatal patterning genes in *Arabidopsis* are indicated with grey dots.

Many of the QTL for both anatomical and functional traits were located in clusters. 24 clusters were identified and named in sequence order (Fig. 7; Table S1; e.g. Chr1A – Chr1D for clusters on chromosome 1 based on their genetic position). The number of QTL in a cluster varied from two (Chr4A, Chr5C, Chr6C, Chr7C, Chr9C, Chr10B) to twelve (Chr6B). There are many examples of QTL co-localizing for traits that are closely related. For example, SCL, SCLWR and SCA in cluster Chr2A or SCD, SCTA, SI and SPI in cluster Chr1B. Interestingly, only two clusters are limited to QTL from a single trait category of stomatal complex size traits, pavement cell traits, stomatal density and index traits or gas exchange traits. Cluster Chr4A contained QTL only for stomatal size traits and cluster Chr9C contained QTL only for pavement cell traits. The other 22 QTL clusters span at least two trait categories (Fig. 7; Table S1). The clusters Chr1C, Chr6A, Chr10A and Chr10B are notable for including overlapping QTL for both epidermal anatomy traits and photosynthetic gas exchange traits.

When QTL were independently identified for the same trait in both years, the direction of the allelic effect was always consistent (Fig. 7; Table S1). Allelic effects were also generally consistent with the trait correlations previously reported. As examples, all allelic effects for QTL at a given locus had opposing directions for SCD versus SCA, or PA versus PD. However, the direction of allelic effects at any individual locus was generally, but not universally, predictable from the trait means of the parental lines. For example, the MS71 allele resulted in lower SCD at 10 of the 17 loci where QTL for SCD were identified, as would be consistent with the lower trait mean for the MS71 inbred line versus B73 (Fig. 7; Table S1). And, the MS71 allele resulted in greater SCA at 7 of the 12 loci where QTL for SCA were identified, as would be consistent with the greater trait mean for the MS71 inbred line versus B73. Consistent with trait values for the parental lines, all of the statistically significant MS71 alleles resulted in lower *g*_s_ relative to B73 alleles. In contrast to other QTL, MS71 alleles in cluster Chr1C were associated with lower gs and greater SD, highlighting the complexity of genetic control of these traits.

## DISCUSSION

Deep-learning has been proposed as a solution for a wide variety of applications in plant phenotyping (Ubbens and Stavness, 2017; Mochida et al., 2018; Singh et al., 2018; Jiang and Li, 2020). Despite this promise and publication of a number of tools, no solution has been widely adopted to assess stomatal patterning. This study successfully met the goals of building, testing, and demonstrating the use of a high-throughput phenotyping pipeline, including automated image analysis by use of machine learning for stomatal patterning traits in a model C_4_ species. This was applied to two-years of samples taken from a field-grown RIL population to advance understanding of the genetic architecture and trait relationships of stomatal patterning and leaf photosynthetic gas exchange in maize. Understanding of genetic variation in stomatal development and function is particularly poor in C_4_ species. As such, the study addresses both technical and biological knowledge gaps that have been long-standing despite the considerable advances in understanding stomatal biology that have been made in recent years (Lawson and Vialet-Chabrand, 2019; Harrison et al., 2020; McKown and Bergmann, 2020).

### High-throughput phenotyping pipeline for stomatal patterning traits

#### Data Acquisition

Optical tomography (OT) was an effective method for imaging the leaf epidermis of diverse maize lines (Fig. 2; Fig. S3). This proof-of-concept built upon previous applications in individual genotypes of *Arabidopsis* (Haus et al., 2018), tobacco (Głowacka et al., 2018) and other dicot species (Haus et al., 2015). Each field of view could be acquired in less than 1 minute, so sampling four or five fields of view per leaf allowed 60 leaves to be comfortably screened with a single microscope in a standard 8-hr work day. This was more efficient and less arduous than our experience of taking leaf impressions or epidermal peels.

Data describing 11 different traits related to stomatal patterning were all significantly correlated across the two growing seasons, despite variation in the growing environment in the field (Fig. 5; Fig. S2). And, this led to consistent findings on trait relationships and the genetic architecture of stomatal traits across the years (Figs. 6, 7, S6).

#### Image Analysis

The Mask R-CNN machine learning tool was successfully trained to automatically locate cells, identify cell classes, segment boundary coordinates and extract density and size traits for stomata as well as pavement cells of maize leaf epidermis. Automatic image analysis was more than 100 times faster than manual measurement of all traits (Table 1). Correlations between the number of stomata and pavement cells identified and counted by the computer versus expert humans were very strong (r^2^ > 0.96) and showed little bias (Fig. 3A,B). This reflected robust predictions across a range of cell morphologies and image qualities, including for partial cells on image edges, and pavement cells above veins (Fig. S7). A second validation step that analyzed all available images for six genotypes that represented the range of SCD and PD in the RIL population suggests the variance is mainly coming from biological replicates, instead of technical errors (Fig. 3C,D). So, the pipeline produced equivalent or higher quality data much more rapidly.

Correlations between computer generated estimates and human assessment of traits describing stomatal complex size were also highly significant (Fig. 4). This aided detection of consistent results across seasons (Fig. 5), and was achieved despite the additional challenge of stomatal size varying less across the RIL population (~50%) than SCD (>100%). Nonetheless, accurate and precise estimation of stomatal size, and SCW in particular, pushed the limits of image resolution when data were collected with the 20X objective lens used in this study. While this approach did allow many QTL and trait relationships to be identified, additional imaging using higher magnification lenses to deliver greater resolution from the OT will likely deliver further gains in phenotyping of these traits.

The pipeline represents a valuable technical advance because previously published automatic stomatal detection and counting algorithms: (1) used data that was collected by slow and laborious methods (e.g. Aono et al., 2019; Bhugra et al., 2019; Sakoda et al., 2019); (2) were limited to detecting stomata and not pavement cells (e.g. Dittberner et al., 2018; Fetter et al., 2019; Li et al., 2019; Sakoda et al., 2019); (3) did not achieve the same accuracy (e.g. Duarte et al., 2017; Saponaro et al., 2017; Bourdais et al., 2019); or (4) were demonstrated to work only within the constrained variation of a limited sample set, which did not include demonstrated applicability for quantitative genetics (e.g. Aono et al., 2019; Fetter et al., 2019; Li et al., 2019). While previous studies achieved these goals individually, combining these features resulted in a tool that could be applied to addressing knowledge gaps about the genetic architecture of SCD and SI in maize.

The independent application of the same tool to stomatal counting in grain sorghum suggests that, with the appropriate training, it has the flexibility and power to be widely applicable (Bheemanahalli et al., in review). But, as with all machine learning solutions to image analysis, there are significant questions about the context specificity of the model used. In the current study, the focus was on development of a method that was robust across a RIL population of a model C_4_ grass species, which included significant variation in many patterning traits but was also subtle relative to large datasets that span many species (Sack et al., 2003). Additional work will be needed to test if new models need to be trained for each individual mapping population or species of interest. One option may be transfer learning methods (Singh et al., 2018) to accelerate the development of machine learning models for new species or even a generic model. Even if this is not possible, training the Mask R-CNN tool required relatively few training instances (33 images containing roughly 2000 cells for stomatal traits and 9000 cells for pavement cell traits). So, building new models for different applications should be a tractable goal.

### Trait variation across the RIL population and years

SCD of maize B72 x MS71 RILs showed a similar range to intraspecific variation in faba bean (Khazaei et al., 2014), wheat (Schoppach et al., 2016; Shahinnia et al., 2016), *Arabidopsis* (Dittberner et al., 2018) and rice (Kulya et al., 2018; Laza et al., 2010). Mean SCD and SCL of the RIL population were very similar to the abaxial trait values for maize and in the mid-range of a diverse set of species previously reported by (McAusland et al., 2016). Therefore, maize does not represent an unusual extreme in terms of epidermal phenotype. Thus, the methods and biological discoveries here may relate to other species. Although, further comparative work is needed as grass epidermal patterning is distinct from that of dicots, and C_4_ species may be expected to differ from C_3_ relatives as a result of broader differences in leaf development and function associated with Kranz anatomy and associated biochemical specialization (Larkin et al., 1997).

The temperature of the 2017 growing season was similar to 2016, but there was ~43 % less precipitation (Fig S2). While this would normally be expected to drive plasticity in stomatal patterning traits, irrigation was applied to avoid plant drought stress in 2017. Consistent genetic variation in stomatal patterning traits between the two years suggests that these traits are, at least, moderately heritable (Fig. 5). SLA differed between years, probably as a result of harvesting material directly from the field in 2016 (low SLA due to high non-structural carbohydrate content) versus after leaves had been held in the lab for photosynthetic gas exchange measurements in 2017 (higher SLA after starch reserves were respired under low light conditions in the laboratory). Nonetheless, genetic variation in SLA was correlated across years and relationships between SLA and other traits were similar across years. Therefore, the resulting data for all traits should be highly amenable for studying trait relationships and QTL mapping. Getting such information under mesic conditions without significant drought stress is valuable because it reduces the likelihood of complex plant-environment interactions that can complicate investigation of genetic variation in *iWUE* and associated traits (Leakey et al., 2019).

### Trait relationships

For the maize B73 x MS71 RIL population, leaf photosynthetic traits and stomatal patterning traits clustered into largely separate groups within which many traits were correlated (Fig. 6). But, there were relatively few correlations between stomatal patterning traits and leaf photosynthetic traits. Most notably, while the classic trade-off between SCD and SCA was observed, there was no significant correlation between SCD or SCA and gs or any other gas exchange trait. This contrasts with the widely held expectation that greater g_s_ will be associated with larger numbers of smaller stomata (Dow et al., 2014; Faralli et al., 2019). This expectation is strongly grounded in theory and data from broad fossil-based comparisons over phylogenetic space and geological time (Franks and Beerling, 2009). Significant relationships between SCD and water fluxes have also been observed in experiments on intraspecific variation in sorghum (Muchow and Sinclair, 1989), rice (Panda et al., 2018), and barley (Miskin et al., 1972). But, there are also a number of studies where SCD was not correlated with *g_s_* in wheat (Liao et al., 2005), rice (Ohsumi et al., 2007), and barley (Jones, 1977). This discordance among studies, and the relatively weak nature of the relationship between SCD and g_s_ that is observed when it does occur within species, indicates how incompletely these structure-function relationships are understood. Therefore, the high-throughput phenotyping methods presented here, which can allow analysis across more and different types of genetic variation, will be valuable. One benefit of testing trait relationships within a RIL population is that the recombination of parental alleles resulting from making crosses breaks up gene linkage that can result from selection and underlie trait relationships, providing a more direct test of the biophysical basis for trait relationships (Des Marais et al., 2013).

It was assumed that the dimensions of stomatal complexes provided information about the maximum size of stomatal pores, based on previous reports for C_4_ grasses (Taylor et al., 2012) and tomato (Fanourakis et al., 2015). Significant correlations were observed between leaf gas exchange traits and SCL, SCW and SCLWR (Fig. 6). Even though there was no relationship between gs and overall SCA, greater gs was associated with stomatal complexes being wider and shorter. This would be consistent with the morphology of the stomatal pore and/or the guard cells and subsidiary cells that surround it playing an important role in determining steady-state gas fluxes (Harrison et al., 2020). And, it suggests that the structure-function relationships of stomatal size-WUE in C_4_ species may parallel those previously reported in *Arabidopsis* (Des Marais et al., 2014; Dittberner et al., 2018). But, the influence of these traits on steady-state gas exchange is much less well understood than its influence on the dynamics of stomatal opening and closing (McAusland et al., 2016). It is also possible that trade-offs between stomatal density, stomatal size and the extent of stomatal opening mean that accurate predictions of gs are possible only when all three of these traits are accounted for. It is also possible that variation in stomatal patterning between abaxial and adaxial leaf surfaces influenced gs in a way that was not captured in the dataset on abaxial traits reported here. But, there are approximately 50% more stomata on the abaxial surface, so it should exert more influence. And, SI of the two leaf surfaces are correlated across diverse maize inbred lines (Michael Mickelbart, pers. comm.).

Understanding the basis for genetic variation in *iWUE* is important because of the benefits to crop productivity, sustainability and resilience that result from improving this key resource use efficiency (Leakey et al., 2019). Greater *iWUE* was strongly associated with lower gs and more weakly associated with lower *A* (Fig. 6). This was consistent with studies on sorghum (Kapanigowda et al., 2013; Fergsuson et al., in review) and switchgrass (Taylor et al., 2016), although the strength of the correlations in maize were significantly stronger. And, it supports the notion that selection for low *g_s_* without equivalently large decreases in *A* may be an approach to improving *iWUE* (Leakey et al., 2019). Of all the stomatal patterning traits, SCLWR had the strongest correlation with *iWUE* (r = 0.28). It meant that longer, narrower stomatal complexes were associated with lower *g*s and greater iWUE (Fig. 6). While this explained only a modest proportion of variation in *iWUE*, it was equivalent to the strength of the relationship between each of the leaf gas exchange traits and SLA, which is widely recognized as a key component of the leaf economic spectrum across broad phylogenetic space (Wright et al., 2004) as well as for C_4_ grasses (Atkinson et al., 2016). SCLWR was not associated with variation in PD, PA or PTA (Fig. 6). This opens up the possibility that this apparently important trait might be manipulated by breeding or biotechnology with minimal unpredictable side effects on epidermal patterning in general. However, the detailed information on epidermal cell allometry provided by the OT images and machine learning algorithm used in this study does also reveal complex relationships among cell types on the leaf surface. For example, PA and SCA are positively correlated, as are SCD and PD (Fig. 6). And, this is consistent with genetic variation in cell size being general in nature across the two major classes of epidermal cells types. However, this occurs at the same time as the tradeoff between SCD and SCA. So, a decrease in SCD appears to coincide with a compensatory increase in PA to fill the available space rather than an increase in PD. And, while SCL and SCW both drive variation in SCA, they are not correlated with each other, and they have opposing relationships with SI, SPI, SLA and the gas exchange traits (Fig. 6). Evaluating how stomatal complex size and proportion varies when SCD is manipulated transgenically may help reveal the key interdependencies between traits.

### QTL mapping

Of 60 QTL identified in 2016 and 83 QTL identified in 2017, 36 were consistently observed in both years (Fig. 7). Additionally, 24 hotspots of overlapping QTLs for multiple traits were identified. The number and strength of QTL identified for leaf gas exchange traits (1-3 QTL per trait in a single experiment) were similar to previous studies of those traits (Hervé et al., 2001; Teng et al., 2004; Pelleschi et al., 2006). In contrast, a greater number of QTL were identified for many of the stomatal patterning traits (e.g. PD – 7, SI – 10, SCA – 10, SCD – 12, SCTA – 7 QTL in a single experiment) than in previous studies (Vaz Patto et al. 2003, Hall et al. 2005, Laza et al. 2010, Schoppach et al. 2016, Shahinnia et al. 2016, Liu et al. 2017, Sumanthi et al. 2018, Delgado et al. 2019; Prakash 2020). This larger number of significant QTL was linked to more small effect QTL (PVE < 10%) being successfully identified. This was unlikely to be the result of false positives because of the consistency in results across the two years of experimentation. This is valuable given the broad evidence suggesting that these stomatal patterning traits are likely to be polygenic, with multiple small effect alleles combining to drive phenotypic variation (Schoppach et al., 2016; Shahinnia et al., 2016; Dittberner et al., 2018; Bheemanahalli et al., in review; Ferguson et al., in review).

Many genes have been implicated in the network regulating cell fate during the differentiation of the epidermis, and therefore stomatal patterning (Pillitteri and Torii, 2012; McKown and Bergmann, 2020). While QTL intervals are too large to allow the causal genes underlying the genotype-phenotype association to be identified, it was possible to determine whether QTL did or did not overlap with the locations of known stomatal developmental genes in maize or orthologs of known stomatal patterning genes in *Arabidopsis* (Table S1). Focusing on the genomic locations where genotype to phenotype associations were identified with greatest overall confidence reveals that orthologs of known stomatal patterning genes were found within the genomic regions of 16 of the 24 QTL clusters identified in this study. For example, an ortholog of EPIDERMAL PATTERNING FACTOR 2 (EPF2, GRMZM2G051168) and Pangloss1 (PAN1, GRMZM5G836190) were co-located within 1 cM of the most significant markers for SCD, PA, *c_i_/c_a_* and *g_s_* in cluster Chr1C (Table S1). PAN1 regulates subsidiary mother cell divisions (Cartwright et al., 2009), while EPF2 is a negative regulator of the number of stomata (Hara et al., 2009). QTL cluster Chr10A co-localized with the maize ortholog of *Arabidopsis* A2-type cyclin CYCA2;1 (GRMZM5G879536). RNAi knock-down of OsCYCA2;1 in rice led to significantly reduced stomatal production, but did not disrupt guard mother cell division, as was the case in *Arabidopsis* (Vanneste et al., 2011; Qu et al., 2018). If confirmed, the involvement of these genes, and others in Table S1, in regulating stomatal patterning in maize would be consistent with the notion that the same set of genes regulates cell fate to control stomatal patterning in dicots and monocots, but the roles of individual genes within the network have been modified over the course of evolutionary time (Raissig et al., 2016; Raissig et al., 2017; Wu et al., 2019). At the same time, the identification of multiple high confidence QTL that do not overlap with existing candidate genes also suggests the possibility that additional genes regulating stomatal patterning remain to be discovered and high-throughput phenotyping of stomatal patterning could aid in their discovery.

The discovery of multiple QTL for many stomatal patterning traits suggests that the goal of reducing *g_s_* and improving *iWUE* by reducing SCD or increasing SWLCR could be achieved through breeding to combine alleles that would result in more extreme trait values than were found in either of the parental inbred lines. This is particularly the case when not all MS71 alleles were associated with, for example, lower SD. Further work is needed to test that possibility and also to determine whether overlapping QTL within clusters are multiple loci in linkage versus the pleiotropic effects of a single locus.

### Conclusion

This study presents an end-to-end pipeline for high-throughput phenotyping of stomatal patterning. New insights were generated on trait relationships within and between stomatal anatomical features and leaf photosynthetic gas exchange. And, the genetic architecture of stomatal patterning and leaf gas exchange traits was characterized in detail. These insights lay the ground work to: (1) apply the high-throughput phenotyping pipeline to other experiments taking quantitative genetics, reverse genetics or forward genetics approaches; and (2) further investigate the physiological and genetic basis for variation in stomatal development, stomatal conductance and *iWUE* in C_4_ species, which is poorly understood despite the agricultural and economic significance of these crops.

## MATERIALS AND METHODS

### Plant material and sampling

Field experiments were done on the University of Illinois at Urbana-Champaign South Farms in Savoy, IL (40°02’N, 88°14’W). Seeds were planted on May 24^th^ in 2016 and May 17^th^ in 2017 with a planting density of 8 plants/m and row spacing of 0.76 m. The crop was grown in rotation with soybean and received 200 kg/ha of nitrogen fertilizer. A population of recombinant inbred lines (RILs) derived from a B73 × MS71 cross was grown, with 197 RILs planted in 2016 and 192 RILs plus the parental lines planted in 2017. This population is a subset of the maize Nested Association Mapping (NAM) population (Yu et al., 2008) and was selected as a result of the parent lines having low (MS71) and moderate (B73) SCD compared to the other inbred founder lines in an initial screen performed at the same field site (Fig. S1). Seeds were obtained from the Maize Genetics Cooperation Stock center (University of Illinois Urbana-Champaign). In 2016, four replicate plants were sampled at random from within the middle portion of nursery rows, which were also self-fertilized for seed production. In 2017, a randomized complete block design was used with two blocks, each containing a replicate plot for each RIL and 6 replicate plots for each parental line. Two sub-samples were collected from separate plants in all replicate rows. In 2017 the field was equipped with drip tape and irrigation was applied uniformly across all genotypes whenever early signs of drought stress were observed. Temperature and precipitation were recorded by the Water and Atmospheric Resources Monitoring Program (Fig. S2). (Illinois Climate Network. 2019. Illinois State Water Survey, 2204 Griffith Drive, Champaign, IL 61820-7495. http://dx.doi.org/10.13012/J8MW2F2Q.)

In both years, measurements were taken on the second leaf beneath the flag leaf following anthesis. In 2016, collection of leaf samples for phenotyping epidermal cell patterning and specific leaf area (SLA) was done in the field. In 2017, tissue sampling was performed after photosynthetic gas exchange measurements were done on the leaves. To allow for this, leaves were cut early in the morning at the base of the leaf blade distally adjacent to the ligule. Cut ends were then submerged in buckets of water and transported to the laboratory. The leaves were then re-cut under water and remained in 50 ml tubes of water during measurements of gas exchange and tissue sampling.

### Epidermal Image acquisition

To phenotype epidermal cell patterning, ~0.5 cm-wide strips were excised from the margin to the mid-rib at a point halfway along the length of a leaf using scissors. Samples were immediately stored in a 2 ml tube, flash frozen in liquid nitrogen, and stored at −20 °C. Leaves were flattened and stabilized onto glass slides with double-sided tape immediately prior to imaging. Abaxial surfaces were imaged with a Nanofocus μsurf Explorer Optical Topometer (Oberhausen, Germany) at 20X magnification with 0.6 numerical aperture. The topography layer was constructed by stacking all the focused pixels across planes of the Z axis. Output images were 0.8mm x 0.8mm on x and y axes (512 x 512 pixels). Five fields of view were scanned on each leaf sample in 2016 and four fields of view were scanned on each leaf sample in 2017. Fields of view were arranged equidistantly along a latitudinal transect from the leaf edge to mid-rib. Sample loss or poor sample quality resulted in incomplete replication for 22 RILs in 2016 and 2 RILs in 2017. Therefore, in total, 3785 images were in the 2016 dataset and 3248 images were in the 2017 dataset (Fig. 1A).

The 3D topographic layer (Fig. 2A) was input into Nanofocus μsurf analysis extended software (Oberhausen, Germany) for image processing as follows: first, non-measured points were filled by a smooth shape calculated from neighboring points. A Robust Gaussian filter with cut-offs of 200μm, 100μm and 100μm were applied in sequence (Fig. 2B). Then, a Laplacian filter with a 13×9 pixel kernel size was implemented (Fig. 2C) before applying another Robust Gaussian filter with a cut-off of 80μm. The final 3D layer was then flattened to 2D in grey scale with auto optimization for luminosity and contrast enhancement.

### Mask R-CNN Model training

Twenty four images were initially randomly selected for training the mask R-CNN model for object instance segmentation. Subsequently, nine additional images of pavement cells that overlie minor veins were added to the training set to improve the detection accuracy for these cells. Each stomatal complex and pavement cell was traced as an object instance using VGG Image Annotator (VIA) (Dutta and Zisserman, 2019). A JavaScript Object Notation (.json) file was generated for each image to record the coordinates for all instance masks within that image. Json files of 26 randomly selected images were pooled to form the training set, and 7 images were pooled into a validation set (i.e. approximately 11,000 unique cells used for model training; Fig. 1A). A Mask R-CNN repository built by Matterport Inc. on GitHub (Waleed, 2017) was used for training on a customized PC with a GeForce GTX 1080 Ti graphics processing unit and 32G of RAM. Model training was based on the ResNet-101 backbone with pretrained weights from the COCO dataset (Lin et al., 2014) with 50 epochs of 100 steps. The learning rate, learning momentum, and weight decay was 0.001, 0.9, and 0.0001, respectively. All images were flipped horizontally and vertically for augmentation. The process taken by Mask R-CNN to make predictions on the instances, size and shape of pavement and stomatal cells is summarized in Fig. 1B.

### Epidermal cell detection, trait extraction and evaluation

The model built during the training process was applied to the detection of cells in the entire image dataset, using the same software and hardware configurations. Instance coordinates and cell type predictions saved by Mask R-CNN model as individual csv files were inputted into R for epidermal trait extraction. The number of stomatal complex and pavement cells within each image were derived as the number of instances detected for these two separate classes and they were standardized by image area to get stomatal complex density (SCD) as well as pavement cell density (PD). The areas of complete, individual stomatal complexes and pavement cells were calculated based on the boundary coordinates using the *splancs* package (version 2.01-40). To derive the stomata complex length (SCL) and width (SCW), an ellipse was first fitted to each stomatal complex using *MyEllipsefit* package (version 0.0.4.2). Stomatal complex width and length were calculated as doubling the radius along the minor and major axis, respectively (Fig. 2G). Total stomatal pore area index (SPI; Sack et al., 2003) is the product of stomatal complex density (SCD) and stomatal complex length (SCL) squared. Stomata index (SI) is the number of stomata divided by the total number of epidermal cells. The *Imager* package (version 0.41.2) and *magick* package (version 2.0) were used to label cells and cell boundaries on detection output images for better visualization.

For validation of SCD and PD, a group of people received training on stomata and pavement cell recognition and reached consensus on the criteria. Two sets of images that were not part of the training dataset were then manually assessed (Fig. 1A). First, six people each manually measured 100 images selected at random from the 2016 and 2017 data. Second, five people each manually measured all images for six genotypes, chosen to represent the range of observed epidermal cell densities, selected from the 2016 dataset. Manual counting was done in Image J 1.8.0 (Schneider et al., 2012) using the multi-point tool. To validate predictions of stomatal size traits by Mask R-CNN, 6 humans each manually measured the same 5 stomatal complexes in each of 42 randomly selected images that were not part of the training dataset (Fig. 1A

### Leaf photosynthetic gas exchange and SLA

In 2017, photosynthesis and stomata conductance were measured using four LI-6400 portable photosynthesis systems incorporating an infrared gas analyser (IRGA) (LI-COR, Lincoln, NE, USA) that were run simultaneously using the protocol of Choquette et al. (2019). 4 leaf disks were sampled using a leaf punch from the same leaf sampled for stomata scanning. Leaf disks were dried in an oven at 60 °C before being weighed on a precision balance (Mettler Toledo XS205, OH, USA). SLA (cm^2^g^−1^) was calculated as the area for leaf punch divided by the mean leaf disk weight.

### Statistical analysis

All statistical analysis was performed in R (version 3.6.0, https://www.r-project.org). Pearson correlations were performed and visualized using *corrplot* package (version 0.84).

The genetic map for B73 x MS71 population consists of 1478 SNPs distributed across all 10 chromosomes of maize (McMullen et al., 2009). SNP data were available as part of the Maize Diversity Project (https://www.panzea.org). Markers were phased and imputed to a density of 1 centiMorgan (cM) resolution. Quantitative trait loci (QTL) mapping for two years was done separately and performed in R for each individual trait using the *stepwiseqtl* function with Haley-Knott algorithm from package *qtl* (Broman et al., 2003) to create a multiple QTL model. A multi-locus model was generated using the stepwise forward selection and backward elimination. The Logarithm of the odds (LOD) penalties for QTL selection were calculated using the *scantwo* function with 1000 permutations for each trait at significance level of 0.05. Following Dupuis and Siegmund (1999) and Banan et al. (2018), 1.5-LOD support intervals were used for each QTL hit. Co-localized QTL were grouped into “clusters” based on their mapping to same or neighboring markers where confidence intervals overlapped. The few QTL with very large confidence intervals (>50 cM), were excluded from clusters. Clusters were named in sequence order (Fig. 7; Table S1; e.g. Chr1A – Chr1D for clusters on chromosome 1 based on their genetic position). Maize 5b gene model coordinates and annotations were both downloaded from MaizeGDB (https://www.maizegdb.org).

## Acknowledgements

This work was supported by a grant from the NSF Plant Genome Research Program (PGR-1238030) and the University of Illinois at Urbana-Champaign Center for Digital Agriculture. Jiayang Xie was supported by a Foundation for Food and Agriculture Research Fellowship. We thank Anthony Studer for helpful discussions on QTL mapping and Elizabeth Ainsworth for comments on a draft manuscript. We thank Patrick Brown, Christopher Montes, Crystal Sorgini and Benjamin Thompson for assistance with acquisition of germplasm, as well as establishment and maintenance of field plots. We thank Timothy Wertin, Nicole Choquette, Jim Berry, Aya Bridgeland and Chris Moller for assistance with sample and data collection. We thank Bindu Edupulapati, Kayla Raflores, Varun Govind and Vishnu Chavva for assistance with manual assessment of stomatal traits in OT images.

---

FIGURE S1. Initial screening of stomatal complex density (SCD; A), pavement cell density (PD; B) and stomatal index (SI; C) for maize NAM founder lines grown in year 2014 (n = 4). Error bars indicate standard errors.

FIGURE S2. Daily mean temperature (red line; °C) and water inputs to field trials (blue bars = total daily precipitation, red bars = irrigation; mm) in Savoy, Illinois for each day of year (DOY) in the 2016 (A) and 2017 (B) growing seasons.

FIGURE S3. Scatterplots of variation among six expert human evaluators in manual measurements of stomatal patterning traits from 100 randomly selected optical tomography images from the B73 x MS71 maize RIL population: stomatal complex density (A), pavement cell density (B), stomatal complex width (C), stomatal complex length (D) and stomatal complex area (E). Data are sorted on the x-axis by rank of the mean trait value for each genotype. The color of a data point corresponds to the human evaluator.

FIGURE S4. Frequency distributions of stomatal complex density (SCD), stomatal complex width (SCW), stomatal complex length (SCL), stomatal complex area (SCA), stomatal complex total area (SCTA), stomatal complex length to width ratio (SCLWR), pavement cell density (PD), pavement cell area (PA), pavement cell total area (PTA), stomatal index (SI), stomatal pore area index (SPI), specific leaf area (SLA), rate of photosynthetic CO_2_ assimilation (*A*), stomatal conductance (*g_s_*), ratio of leaf intercellular to atmospheric CO_2_ concentration (*c_i_/c_a_*) and intrinsic water use efficiency (*iWUE*) for the maize B73 x MS71 RIL population in grown in 2016 (grey) and 2017 (yellow). The mean trait values from 2017 for the parent lines MS71 (orange) and B73 (blue) are plotted.

FIGURE S5. Correlation matrix for stomatal complex density (SCD), stomatal complex width (SCW), stomatal complex length (SCL), stomatal complex area (SCA), stomatal complex total area (SCTA), stomatal complex length to width ratio (SCLWR), pavement cell density (PD), pavement cell area (PA), pavement cell total area (PTA), stomatal index (SI), stomatal pore area index (SPI), specific leaf area (SLA), based on genotype means of the maize B73 x MS71 RIL population grown in 2016 (n = 197). Statistically significant correlations (p<0.05) are highlighted with colored cells that reflect the strength of the correlation by the size of the shaded area and are colored from red (positive correlation, coefficient = 1) to blue (negative correlation, coefficient = −1).

FIGURE S6. Sum of percentage of variance explained (PVE) for all QTLs identified for each trait in 2016 (grey bars) and 2017 (yellow bars). Traits are presented in rank order from greatest to least sum PVE: stomatal index (SI), stomatal complex area (SCA), stomatal complex density (SCD), stomatal complex total area (SCTA), pavement cell area (PA), pavement cell density (PD), stomatal complex width (SCW), stomatal pore area index (SPI), stomatal complex length to width ratio (SCLWR), pavement cell total area (PTA), specific leaf area (SLA), stomatal complex length (SCL), stomatal conductance (*g_s_*), ratio of leaf intercellular to atmospheric CO_2_ concentration (*c_i_/c_a_*), intrinsic water use efficiency (*iWUE*), rate of photosynthetic CO_2_ assimilation (*A*). Gas exchange traits were only assessed in 2017.

FIGURE S7. Examples of input images and the predictions of cell instances made for them across a range of epidermis morphology and image qualities, including: pavement cells above veins (where veins are highlighted with arrows; A, B); lower quality images (C, D), and a darker image (E).

